# Updated phylogeny of Chikungunya virus suggests lineage-specific RNA architecture

**DOI:** 10.1101/698522

**Authors:** Adriano de Bernardi Schneider, Roman Ochsenreiter, Reilly Hostager, Ivo L. Hofacker, Daniel Janies, Michael T. Wolfinger

## Abstract

Chikungunya virus (CHIKV), a mosquito-borne alphavirus of the family Togaviridae, has recently emerged in the Americas from lineages from two continents, Asia and Africa. Historically, CHIKV circulated as at least four lineages worldwide with both enzootic and epidemic transmission cycles. To understand the recent patterns of emergence and the current status of the CHIKV spread, updated analyses of the viral genetic data and metadata are needed. Here, we performed phylogenetic and comparative genomics screens of CHIKV genomes, taking advantage of the public availability of many recently sequenced isolates. Based on these new data and analyses, we derive a revised phylogeny from nucleotide sequences in coding regions. Using this phylogeny, we uncover the presence of several distinct lineages in Africa that were previously considered a single one. In parallel, we performed thermodynamic modeling of CHIKV untranslated regions (UTRs), which revealed evolutionarily conserved structured and unstructured RNA elements in the 3’UTR. We provide evidence for duplication events in recently emerged American isolates of the Asian CHIKV lineage and propose the existence of a flexible 3’UTR architecture among different CHIKV lineages.

## 1. Introduction

Phylogeographic inquiries can reveal viral transmission patterns and pathogenesis of viral diseases. Dramatic (re)emergences of arthropod-borne viruses (arboviruses), such as Chikungunya (CHIKV), West Nile, Zika, and Dengue have increased in frequency and intensity, thus raising public health concerns [1]. Prolific expansion of arboviruses in countries where the viruses have previously not been recorded has drawn public attention. These trends are considered to be a function of climate change effects on the geographic range in which the vector can complete its life cycle and transmit CHIKV [2, 3]. Here, we investigate the RNA architecture of CHIKV untranslated regions (UTRs) that underlies their ability to adapt and transmit in new environments and provide an updated large-scale phylogeny summarizing the geographic spread of CHIKV lineages.

CHIKV is a mosquito-borne virus that belongs to the *Alphavirus* genus, which encompasses mainly arboviruses, and is believed to have a marine origin [4, 5]. The presence of CHIKV in sylvatic mosquitoes, such as *Aedes africanus, Aedes furcifer* and *A. taylori*, and non-human primates in Uganda and Tanzania suggests that the virus originated in Central or East Africa [6, 7].

The first isolation of CHIKV was in 1952 in the Newala district of Tanganyika, a British territory that is now part of Tanzania [8, 9]. CHIKV has caused multiple outbreaks in all continents except Antarctica, most recently in Europe and Central and South America. In total, CHIKV has been present in over 100 countries [6, 10–12].

CHIKV infects multiple human tissues, causing a febrile illness which is typically characterized by myalgia, polyarthralgia, polyarthritis, and rashes [13]. Several vaccine candidates are currently in preclinical or clinical development [14].

The virus had not been associated with life-threatening symptoms before an outbreak in 2005 on La Réunion island off the East African coast [15], which demonstrated a drastic expansion of the disease spectrum. In La Réunion and after, higher morbidity has been associated with CHIKV infection, including neurological issues, such as visual and hearing loss, paralysis, Guillain Barré Syndrome (GBS) and renal complications [13, 16, 17]. Recent mutations that change protein structure have been attributed to increased ability to establish infection (infectivity) and outbreaks as the virus becomes better adapted to vector species (e.g., *A. albopictus*, the only suitable agent for transmission in La Réunion island) [18]. Species of the *Aedes* genus, in particular *A. albopictus*, *A. aegypti*, *and A. polynesiensis*, are carriers of CHIKV, capable of transmission to humans [19].

### 1.1. Chikungunya lineages

CHIKV is a virus for which four major lineages have been proposed: Asian Urban (AUL), Indian Ocean (IOL), East, Central and South African (ECSA), and West African (WA) [20, 21]. Despite these notions, the origin and the history of how CHIKV spreads inside of Africa remains elusive. This gap in our understanding is due to undersampling and confounding factors such as co-circulating Dengue and Yellow Fever viruses. The co-circulation of viruses diminishes the value of serology test which are not species specific. While the AUL has historically been constrained in Southeast Asia, there have been recent outbreaks in the Americas [18]. The IOL, circulating in India and surrounding regions, has been mainly restricted to cause outbreaks in South and Southeastern Asia, although it has caused recent outbreaks in Europe. The WA lineage is entirely isolated.

The ECSA lineage has a unique history as it gave rise to the IOL as well as has also been identified as the origin of recent outbreaks in Brazil and Haiti [12, 22, 23]. The vast territory in Africa, the lack of sampling, and increasing diversity of ECSA lineage fosters doubt on whether ECSA is a single lineage.

### 1.2. Conserved RNA structure as evolutionary trait

The CHIKV genome consists of a single-stranded, (+)-sense RNA of approximately 11.8kb, encoding nine proteins which are divided into structural and non-structural proteins [24]. The genome is organized as two open reading frames (ORFs), flanked by structured UTRs [25].

Populations of RNA viruses exhibit large genetic variability. This variation allows populations to have variants that are suited to changing conditions and immune escape. Viral variation results mainly from two mechanisms, mutation and recombination. Mutation occurs during replication by mis-introduction of nucleotides by error-prone RNA-dependent RNA polymerases that lack proof-reading capacity. Recombination occurs when co-circulating lineages co-infect a host and mixed segments are incorporated into the descendent viruses [26].

The role of functional RNA elements in pathogenesis is well established in flaviviruses [27, 28], where they are typically conserved evolutionarily in UTRs [29–31]. In CHIKV, patterns of coupled historic recombination and mutation events are encoded in the 3’UTRs of different lineages [32]. The resulting variants have highly divergent 3’UTRs that vary in length. Characteristics of these 3’UTR variants are variable number of direct repeats (DRs) at the sequence level [2, 33]. Distinctive CHIKV lineages also differ in their 3’UTR architecture, in particular in the copy number and arrangement of sequence repeats [34].

Evolutionary conservation in RNA viruses is typically maintained in secondary structure and manifested in rich base-pair conserving covariation patterns [35]. Relatively little is known about structured RNAs in alphaviruses, both in coding and in non-coding regions. Coding regions evolve under strong selection pressure to maintain protein function. Nevertheless, it has become apparent that structure within coding regions can also have biological function, as seen in the HIV RRE element [36]. In the context of alphaviruses, Kutchko *et al.* [37] performed RNA structure probing experiments, focusing on Sindbis virus (SINV) and Venezuelan equine encephalitis virus (VEEV), but could not find evidence for broad conservation of functional RNA structures.

The knowledge of structured RNAs in alphavirus UTRs, and CHIKV in particular, is limited. This limitation results from the fact that earlier studies focus on lineage-specific sequence repeats and could not reliably establish a relationship between all observed repeat patterns and RNA secondary structure [32, 34].

A large number of genomic sequences and metadata have been deposited recently to public repositories. These data have raised the possibility of revising the lineages of CHIKV as a whole, reevaluating the historical global spread of the major epidemic clades and evaluating the RNA structural patterns among different clades. We also set out to structurally characterize the UTRs of all known CHIKV lineages, and found evidence for lineage-specific structured and unstructured repeat elements. We hypothesize such RNA elements to be involved in viral replication, vector adaptation and specificity, host factor binding, and pathogenicity.

## 2. Materials and Methods

### 2.1. Taxon sampling

We obtained viral genome data from the National Center for Biotechnology Information (NCBI) `Genbank` database (https://www.ncbi.nlm.nih.gov/genbank/). We downloaded 598 whole genome sequences of CHIKV. We compiled all geographic metadata available in these genome records to create data sets for the analyses described below. We removed eight sequences from the data set due to missing geographic location metadata or designation as a vaccine sequence. The metadata associated with geographic location was associated with the United Nations geoscheme for consistency. Therefore, “Central Africa” is labeled “Middle Africa”.

### 2.2. Phylogenetic tree search

Two complete ORFs of CHIKV were extracted from the complete viral sequences for the phylogenetic analysis. Multiple sequence alignments of the individual ORFs were computed using `MAFFT` [38] under default settings and visualized in `AliView` [39]. In order to examine the possibility of recombinant sequences, we used RDP and methods therein (RDP, GENECONV, Bootscan, MaxChi, Chimaera, SiScan and PhylPro)[40].

We conducted phylogenetic tree searches with partitioning to separate the two ORFs. We used tree search method under the optimality criterion of maximum likelihood with substitution model testing as implemented in `IQ-Tree` [41].

We executed a 1000-replicate ultrafast bootstrap analysis of clade frequencies [42] and a SH-aLRT measure of support [43]. The ultrafast bootstrap analysis is a variation of the traditional bootstrap using heuristics and constraints to speed the search for optimal tree topologies. The SH-aLRT is an approximation of the likelihood ratio, a direct measure of how much the evidence supports the hypothesis.

According to recently published phylogenies of CHIKV [20, 22, 44], viruses from the Western African clade form a sister group to all other CHIKV clades. Given these results and the absence of co-circulating Western African strains with any other CHIKV lineage, we selected this clade as the outgroup for the phylogeny of CHIKV. de Bernardi Schneider [44] observed Western Africa as a monophyletic clade sister to all other clades by utilizing an external outgroup (O’nyong nyong virus), thus the selection of this group should not yield an inherent bias on the tree topology at the same time that it results in a cleaner molecular sequence alignment given the outgroup sequences to belong to the same virus species as the ingroup. We performed the same procedure for a subset containing the 111 complete 3’UTR sequences from the 590 whole genome sequence data set.

The final trees with metadata annotation were rendered using custom Python scripts from the `ETE3` framework [45].

### 2.3. Global spread of virus

We extracted the list of taxa of the three major epidemic clades, Asian Urban Americas (AUL-Am), ECSA-IOL, and ECSA-Middle African South American (ECSA-MASA), from the phylogenetic tree generated from all CHIKV sequences. We created a csv table including all locations from the dataset with corresponding latitude and longitude extracted from getlatlong.net. We performed individual alignments for each epidemic clade on concatenated ORFs and ran these on an R script pipeline (described here). Our script performs five steps:

1. Tree search under the neighbor-joining algorithm with 100 bootstrap replicates [46, 47];
2. Collapsing of branches with a bootstrap value under 80% [48];
3. Replacing tree tip names with metadata information (i.e. geographic location);
4. Parsimony ancestry reconstruction using the *asr_max_parsimony* function from Castor R Package [49, 50] to build an edge list (i.e. origin to destination geographic state transitions);
5. Render the geographic state transitions reconstructed on the previous step on a map using geographic coordinates for each location [51, 52].

In order to evaluate the historical spread of CHIKV using a parsimonious ancestral reconstruction, the tree must be rooted to infer event directionality. Thus, we selected a root for each clade based on the complete CHIKV phylogeny. For each individual clade, the sister taxon to all other taxa was selected. When a clade had multiple sister taxa, we randomly selected one as the root. We selected HM045788.1 (India) as the outgroup for AUL-Am, KF283986.1 (Comoros) for ECSA-IOL and KX262996.1 (Cameroon) for ECSA-MASA.

This script is an adaptation from `StrainHub` (https://github.com/abschneider/StrainHub) [49], a web-based application to reconstruct pathogen transmission networks, to visualize the spread of the individual lineages/epidemic clades. The scripts and data sets along with a detailed explanation of the functions and packages used for the analyses described here are available online at https://github.com/abschneider/paper-chikungunya-phylogeny.

### 2.4. Conserved RNA structures in CHIKV 3’UTR

Evolutionary conservation of functional RNAs is typically achieved by nature at the level of secondary structures. Comparative genomics approaches aim at detecting these conserved, structured RNAs as patterns of covariation in genome sequence alignments, typically constructed from phylogenetically narrow subgroups. To this end, Covariance Models (CMs) are statistical models of RNA structure formation that extend traditional Hidden-Markov-Models (HMMs) by simultaneously representing sequence and secondary structure information [53]. CMs provide a straightforward approach to find homologous RNAs in large RNA sequence databases.

All CHIKV 3’UTRs were processed and curated from full genomes by custom Perl scripts based on the `ViennaNGS` toolbox [54]. Structural RNA alignments were computed with `mlocarna` [55], and consensus structures were generated with `RNAalifold` [56]. Locally stable secondary structures were computed using `RNALalifold` from the `ViennaRNA` package [57]. Homologies of both sequences and secondary structures were inferred by HMMs and CMs as implemented in the `Infernal` package [58] as outlined previously [35]. All secondary structure plots were rendered using the `RNAplot` utility [57].

## 3. Results

### 3.1. Phylogeny of Chikungunya indicates multiple lineages in Africa

No significant recombination events (P-value of < 0.05) were detected in the aligned data sets, therefore we kept all 590 sequences in the data sets. The phylogenetic tree search performed with `IQ-Tree` resulted in a best-scoring heuristic maximum likelihood tree for CHIKV with 590 genomic sequences (Figure 1). Deep ancestral branches that define the lineages had high (>90) SH-aLRT support/ultrafast bootstrap values. More recent branches had lower SH-aLRT support/ultrafast bootstrap values. The phylogenetic hypothesis for CHIKV with branch values is available as supplementary dataset SD1 at https://github.com/abschneider/paper-chikungunya-phylogeny

**Figure 1.**
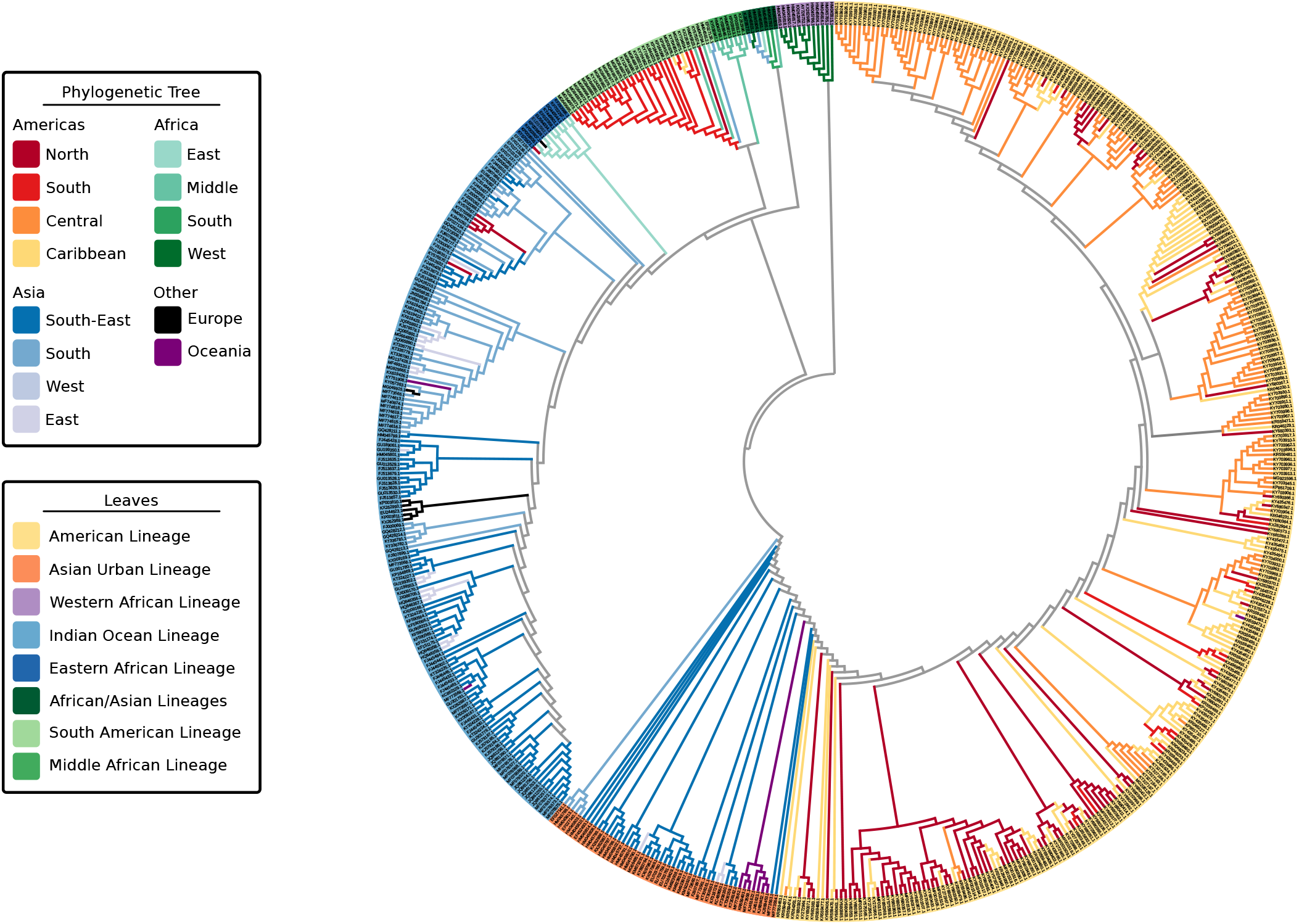
Maximum-Likelihood phylogenetic tree of 590 Chikungunya virus genomic sequences highlighting the geographic locations (tree branches) and lineages (tree leaves). Tree was computed using `MAFFT` alignment of individual ORFs and partitioned tree search with model test using IQ-Tree. Leaves are colored by lineage association. Branches of the tree are colored by geographic location of a sample. Colors of ancestral nodes and branches do not imply geographic origin and therefore are depicted in gray.

The four viral strains grouped as expected according to reports in literature [20, 22], however the increased number of taxa in the tree allowed us to better investigate relationships of clades and infer source-sink events (Figure 2). In addition to the four major lineages AUL, ECSA, IOL and WA, our decision to split the concept of ECSA into its three regions, Eastern Africa, Central (Middle) Africa, and Southern Africa, rose from the presence of distinct clades in this large-scale phylogeny. The epidemic strain from the ECSA-MASA clade was first found in Middle Africa and later identified in South America. The AUL lineage gave rise to an extensively nested American lineage which spans North, South, and Central America, as well as the Caribbean.

**Figure 2.**
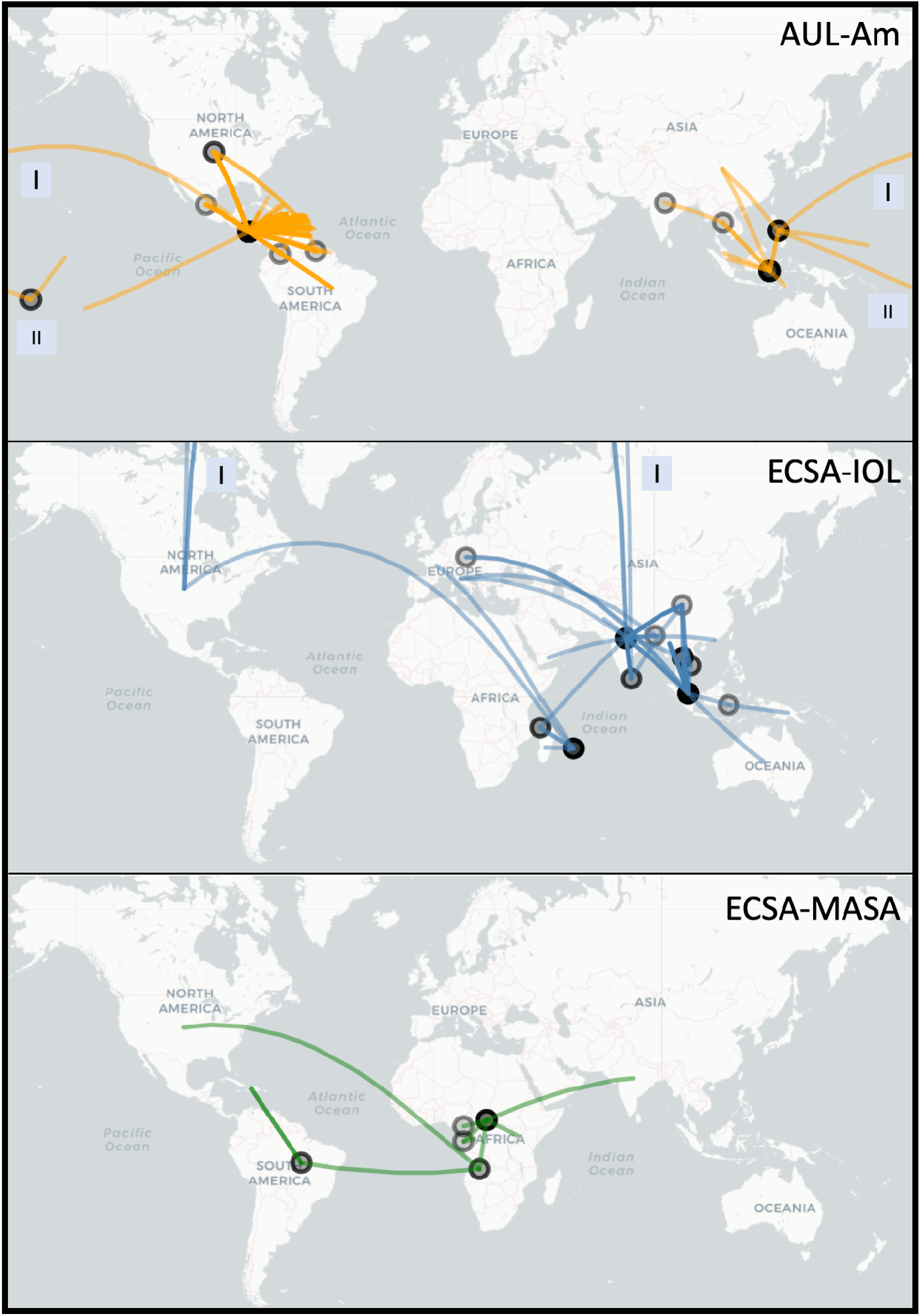
Worldwide spread of major epidemic Chikungunya clades: Asian Urban American (AUL-Am), ECSA Indian Ocean Lineage (ECSA-IOL) and ECSA Middle Africa South America (ECSA-MASA). This figure is rendered using the R package Leaflet with the underlying data based on parsimony ancestral reconstruction of metadata for place of isolation on a the tree resulting from neighbor-joining tree search with collapsed branches under 80% bootstrap threshold (see methods for more details). The circles on map represent locations from which a transmission event originates. The color transparency represents the number of times a transition between characters occur on a location (node) or between locations (edge) (darker lines = more transitions). Boxes labeled “I” and “II” indicate that line crossed the International Date Line (e.g. AUL) or the shortest physical path between the two points (for mapping purposes only) was through the North Pole (e.g. ECSA-IOL).

Thus, we extrapolate the names for these discrete lineages based on geographic location: Indian Ocean, Eastern African, South American, Middle African, African/Asian, Asian Urban, American and Western African (denoted as “leaves,” Figure 1). Some of these lineages belong to the three major epidemic clades: AUL-Am (Asian Urban + American lineages), ECSA-MASA (Middle Africa / South American Lineage) and ECSA-IOL (Eastern Africa + Indian Ocean Lineage). In this paper, we distinguish the individual lineages, which are delimited by geographic region (e.g., IOL, WA), from epidemic clades, which may encompass more than a single lineage (e.g., AUL, ECSA-IOL, ECSA-MASA). Individual limited outbreaks (e.g., IOL in Europe) are not considered new lineages as the outbreak did not generate a new endemic region for the CHIKV lineage (Table 1).

**Table 1.** Comparison between literature and proposed nomenclature of Chikungunya lineages and epidemic clades based on updated phylogeny.

| Previous Work | This Study |  |
| --- | --- | --- |
| Lineages | Lineages | Epidemic Clades |
| Indian Ocean (IOL) | Indian Ocean | ECSA-IOL |
| East Central and South African (ECSA) | Eastern African |  |
|  | South American | ECSA-MASA |
|  | Middle African |  |
|  | African/Asian | N/A |
| Asian Urban (AUL) | Asian Urban | Asian Urban American (AUL-Am) |
|  | American |  |
| West African (WA) | Western African | Western African (WA) |

### 3.2. Spread of major Chikungunya epidemic lineages

Our reconstruction of the spread of the three distinct epidemic clades is in agreement with the current literature (Figure 2). The Asian Urban lineage has spread from Southeastern Asia to the Pacific Islands and Central America, with traveler cases reaching the Continental United States (AUL-Am clade) [7, 59]. A strain from Eastern Africa spread to India and is currently endemic in South and Southeast Asia (ECSA-IOL clade) [60]. More recently, a lineage from Middle Africa was introduced in South America (ECSA-MASA clade) [61, 62].

### 3.3. Lineage-specific 3’UTR organization

Composition and length of the full 3’UTRs analyzed in this study differ substantially between lineages and range from 510 to 930 nucleotides. Earlier studies reported cascades of repeated sequence elements (RSEs) [33] and a structurally conserved sequence element (CSE) required for replication [2] in CHIKV 3’UTR. Chen *et al.* [34] proposed distinct direct repeat elements 1 and 2 (DR1, DR2), together with a structured Y-shaped element, which all appear in lineage-specific duplication patterns. Filomatori *et al.* [32] adopted this classification in a recent study, however they were not able to see *in silico* secondary structure signals in DR sequences except for a Y-shaped element. Here, we follow up on previous knowledge, using the full set of presently available sequence data (i.e. 196 complete CHIKV 3’UTRs), and conclusively infer hitherto unresolved structural properties of all conserved elements.

From a multiple sequence alignment of all available 3’UTRs from all lineages, we manually characterized five highly conserved regions into four structured and one non-structured elements (Figure 3). We inferred sequence and structural homology from searches in the complete UTR set of all lineages, thus characterizing all instances of each conserved element. We used structural alignment and consensus structure prediction to find the conserved structures of the four structured elements.

**Figure 3.**
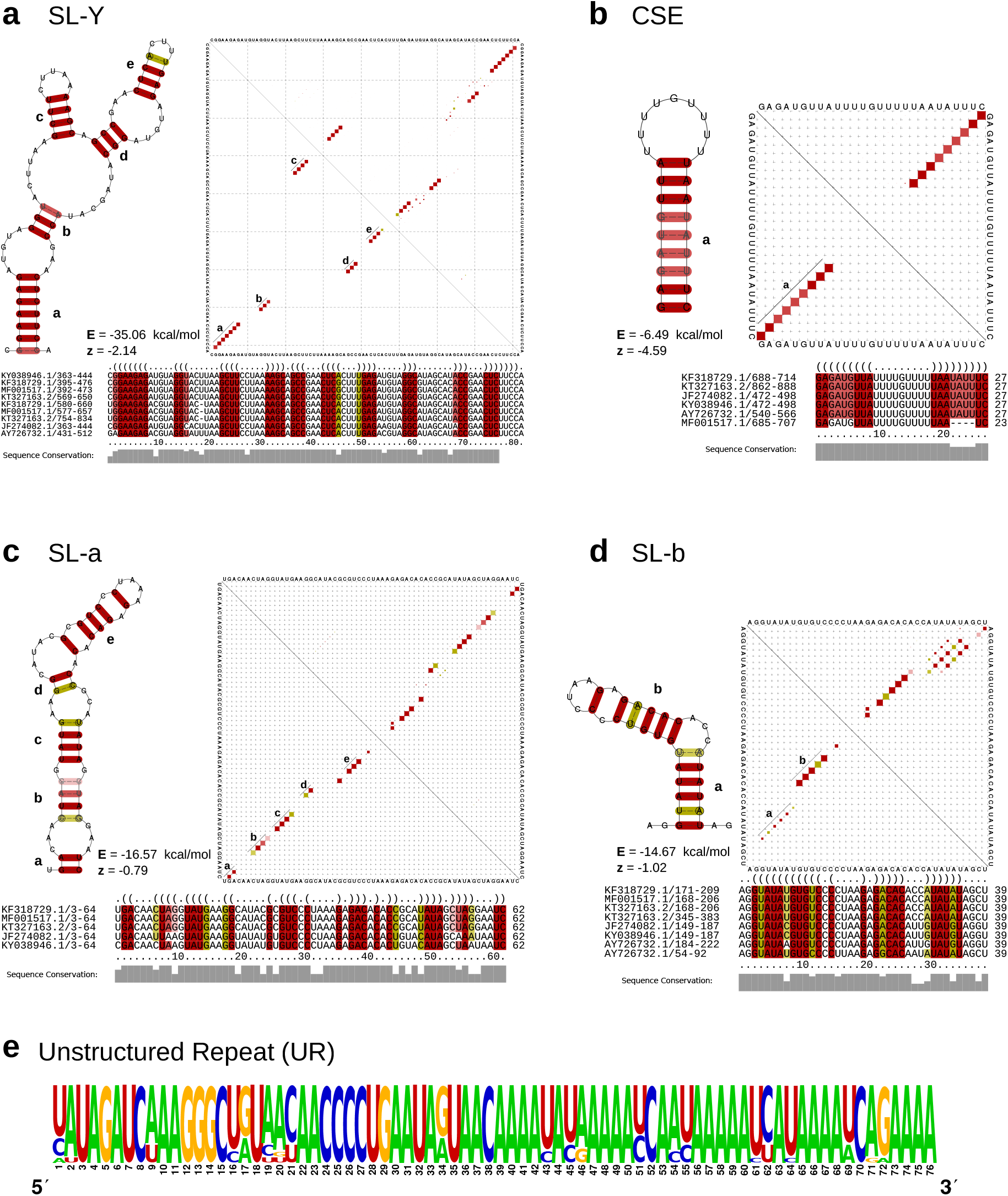
**a-d** Secondary structure cartoons, structural alignments and base-pair probability plots (dot-plots) of the structured CHIKV 3’UTR elements SL-Y, CSE, SL-a, and SL-b. Colored squares in the dot-plot report a pairing of two bases, with full squares corresponding to 100% pairing probability. The base pairings pattern of the corresponding most stable, minimum free energy structures are shown in the lower triangles, while the upper triangles show the base pairing probabilities considering all possible alternative, suboptimal structures. Colors in structure drawings, alignments and dot-plots indicate how many different types of base combinations are found for a particular base pair (red: 1; yellow: 2). All structure plots show the consensus structures as calculated by `RNAalifold` [56]. For each structure, the free energy (E) and the `RNAz` [64] z-score (z) are provided. Negative z-scores indicate a structure is more stable than it would be expected for other RNAs of the same dinucleotide content. **e** Sequence logo of the intrinsically unstructured repeat element UR, computed from all HMM-derived homologous sequences within the data set. Structure cartoons are not shown since the overall base pairing probability is essentially zero for this sequence.

We characterized two stem-loop (SL) elements, termed SL-a and SL-b, the latter being conserved among all lineages (Figure 4). While SL-a is only conserved among the ECSA-derived and AUL lineages, comparison of the two SL elements indicates that SL-b is likely a shortened variant of SL-a and comprised of its innermost hairpin (Figure 3, c, d). Interestingly, this characteristic can also be observed in O’Nyong-Nyong virus (ONNV), which shares common ancestry with CHIKV. The genomic coordinates of both elements approximately correspond to the previously reported DRs, with DR1a spanning over the downstream portion of SL-a and DR1b being consistent with SL-b. The Y-shaped element (SL-Y) is conserved among all lineages but varies in copy number between ECSA-derived/WA (one copy) and AUL (two copies), and shows strong sequence conservation (Figure 3, a). The previously described CSE [63] folds into a short hairpin loop and is found upstream of the poly-A tail in all CHKIV isolates (Figure 3, b).

**Figure 4.**
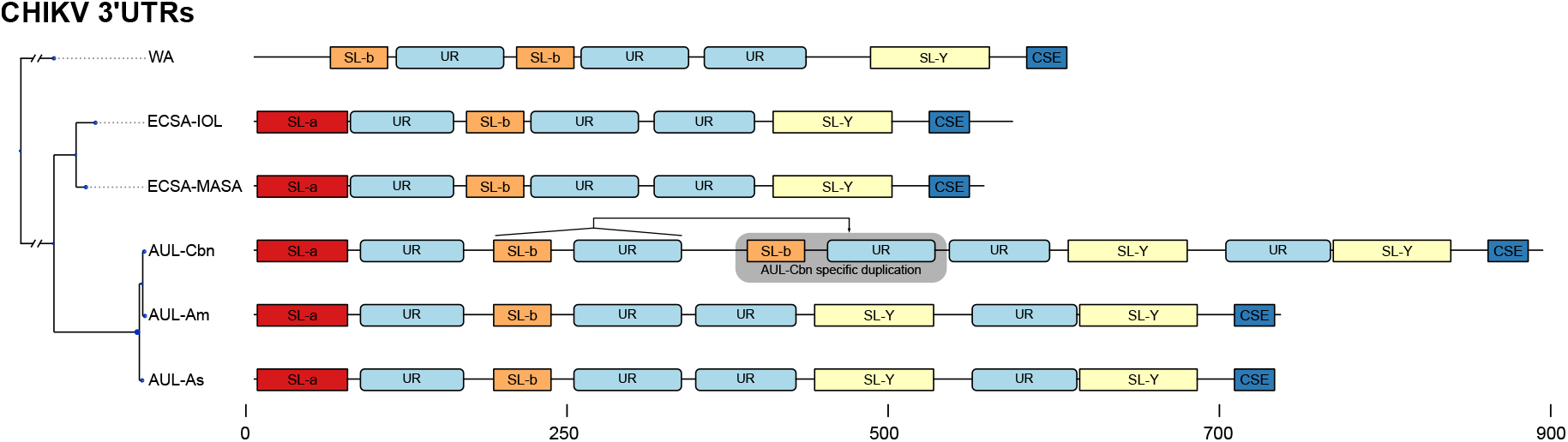
Schematic of the 3’UTR architecture of all major CHIKV strains. The phylogeny on the left is derived from a Maximum-Likelihood tree based on a genomic alignment of six representative sequences. For each strain, the sequence with the longest UTR was selected as a representative (WA: AY726732.1, ECSA-IOL: JF274082.1, ECSA-MASA: KY038946.1, AUL-Cbn: KT327163.2 (Mexico), AUL-Am: MF001517.1 (USA), AUL-Asia: KF318729.1 (China). All boxes correspond to exact coordinates of annotated structural/non-structural elements. A scale bar is provided for the nucleotide coordinates.

The structured elements SL-a, SL-b, and SL-Y are separated by one or two repeats of an intrinsically unstructured repeat (UR) element (Figure 3, e). The total copy number of UR elements varies from three to five between lineages (Figure 4). This sequence repeat element lacks structure, as marked by its incapability to fold into any stable RNA secondary structure. Both single sequence predictions of individual repeats, as well as consensus structure prediction using an alignment of all instances did not yield a stable minimum free energy structure, or alternative base pairing probabilities.

Annotation of all available 3’UTR sequences with CMs and HMMs revealed that different CHIKV lineages have evolved distinct patterns of structured and unstructured functional RNA elements. This lineage-specificity of potentially functional RNAs has led to a varied 3’UTR architecture among CHIKV strains. This is particularly pronounced in the AUL and its descendants, which show a characteristic 3’UTR organization. Several geographically diverse isolates of the AUL-Am (also referenced as Caribbean lineage, AUL-Cbn) appear to have undergone a partial duplication event in the 3’UTR [65]. As a result, an additional copy of the non-structural sequence repeat UR and the structural element SL-b are found immediately downstream to the first copy of SL-b and UR (Figure 4).

## 4. Discussion

Studies on CHIKV phylogenetics have provided phylogenomic means of understanding disease spread in the context of correlating disease epidemiology. The phylogenetic analyses we present are in agreement with the current literature, although the increased number of sequences presented provide insights on the hidden history of ECSA. Interestingly, the phylogeny based on ORFs is in agreement with the phylogeny based on complete UTRs, despite the intuitive idea that different evolutionary forces should act on coding and non-coding regions.

### 4.1. Chikungunya virus nomenclature

It is critical to emphasize the significance of the definition of “lineage” and “epidemic clade” when discussing the evolution of a virus. Much has been proposed as to how species should be delineated. In 1977, Wiley [66] proposed that the definition of species should be that “A species is a single lineage of ancestral descendant populations of organisms which maintains its identity from other such lineages and which has its own evolutionary tendencies and historical fate”. Peterson [67] criticizes the current system of viral taxonomy, mentioning that one criterion alone should dominate viral classification: the evolutionary independence of evolving lineages.

Assessing each CHIKV taxon as an individual, and bearing in mind the concept of phylogenetic independence, we cannot conclusively state that CHIKV sequences isolated in the Americas are divergent enough from ancestral sequences in Africa and Asia to be considered two new American lineages. This logic however, is limited when considering the similar phylogenetic structure of IOL rising from Eastern African taxa. This non-uniformity represents the difficulty in qualifying lineages based on subjective interpretation.

Further, IOL and AUL have region-specific derived clades circulating newly endemic regions. Peterson [67] warns against the use of endemic regions to classify species. We acknowledge the flaw in solely using geography as a naming mechanism as the increased movement between continents due to airport passenger can substantially increase the number of co-circulating lineages named based on geography.

In order to remain consistent with the current literature we kept the naming scheme based on geography, but defined lineage as the pool of individual taxa found in a clade that are not isolated geographically. We define epidemic clade as a monophyletic group that may encompass one or more lineages regardless of geography.

### 4.2. Lineage diversity in Chikungunya virus

Multiple waves of CHIKV infection have occurred over the past two decades, with outbreaks traced from Eastern and Middle Africa to Southeastern Asia (e.g., Thailand, Malaysia), Southern Asia (India) and the Americas [20]. Clades of CHIKV observed in the analyses are consistent with those found in the literature with an important exception. Similar to White *et al.* [23], who alluded to the presence of ECSA subgroups, we observed that ECSA has multiple co-circulating lineages which gave rise to the current outbreaks in Southern and Southeastern Asia and the Americas. With the large amount of new sequences we analyzed, what was once called ECSA, East Central South African lineage, is no longer a single lineage because it is not monophyletic. Figure 1 demonstrates that Middle African taxa gave rise to the outbreak in South America, while Eastern African strains gave origin to IOL. IOL has remained nearly exclusively on the Asian continent, with exception of a few African and European outbreaks. AUL, a sister clade to all ECSA derived lineages, has reached the American continent with extensive nesting diversity (Figure 1). In Africa, AUL is known to have emerged in 2004 and spread explosively in the Comoros islands and incited a severe epidemic in Kenya [18]. In the Americas, strains from AUL are observed as coming from the US to Central America (Figure 2), given the absence of locally acquired cases in the US, the indication of an ancestral sequence originating a transmission event to Central America could be due to lack of sampling the ancestral strain sequence and the close similarity of it with strain sequences found in Central America.

Asian Urban and IOL are the main lineages of CHIKV in Asia [68, 69]. Moreover, Asian Urban has since emerged in Central America in 2013 and South America in 2014[70] forming the AUL-Am clade. The Americas faced a devastating outbreak in 2013-2014 after the AUL lineage was introduced to the Caribbean [71], causing nearly 3 million estimated cases. In 2014, the ECSA lineage was detected in Brazil as well [70]. Based on our phylogeny, the additional sister group lineage to both ECSA-IOL and ECSA-MASA, which encompasses Southern African isolates and is related to a few Asian isolates (of the African/Asian lineage) has no major outbreaks reported.

We propose that the ECSA genotype is too reductive in light of the rapid expansion and increased taxonomic sampling, and should be separated into their respective regions of the African continent: Eastern, Central, and Southern. Nevertheless, it is important to raise the attention to the lack of sampling in Africa which obscures the Chikungunya virus dynamics between the distinct regions within Africa. The current but limited amount of African sequences utilized in this study indicate that there may be multiple co-circulating lineages and this results should serve as justification for the importance of understanding the virus dynamics in Africa. As more isolates are sequenced, a more nuanced history of CHIKV evolution will emerge which will aid in our understanding of the dynamics of these outbreaks.

### 4.3. Lineage-specific 3’UTR architecture

Earlier work in alphavirus 3’UTRs reported a varied architecture of repeat sequence elements (RSEs) [2, 33, 72], which appear to influence virus replication in mosquito cells without affecting infectivity in mammalian cells [34]. Interestingly, molecular evolution experiments with engineered CHIKV 3’UTRs in mosquito and mammalian cells suggest that improved fitness was acquired mainly due to mutations in protein coding regions [73]. Still, the specific biological functions of individual 3’UTR RSEs have not been studied in detail. The comparative genomics approach employed in this study confirms the previously reported repeat organization in CHIKV 3’UTR sequences. It is now possible to conclusively associate repeat sequences with distinct RNA secondary structures that are conserved among all lineages. Importantly, the organization of all CHIKV 3’UTRs follows a common pattern characterized by an alternating arrangement of structured, i.e. SL-a, SL-b and SL-Y, and unstructured regions. The latter are defined by a lack of any RNA structure, as predicted by single sequence and consensus folding algorithms. This suggests that these regions of the CHIKV genome exist in a predominantly unpaired state, rendering them highly accessible.

We hypothesize that such intrinsically unstructured repeat regions act as structural insulators that separate functional secondary structures from each other. Unstructured stretches can mediate the formation of structure in adjacent regions by prohibiting base pairs that are in conflict with the element structure. Likewise, single-stranded, unstructured RNAs can serve as easily accessible protein binding sites. In particular, the enrichment of `A` nucleotides in the unstructured region not only indicates the lack of secondary structure, but might also facilitate the sponging of RNA-binding proteins [74]. Earlier studies suggested an involvement of alphavirus 3’UTRs with cellular HuR and La proteins [75, 76]. While HuR binding in CHIKV appears impaired [77], stabilization of viral RNA through host factors is an evolutionary trait to counteract viral genome degradation by host nuclease activity [25]. Conversely, other RNA-binding proteins, such as the Musashi family of translational regulators have been shown to efficiently interact with viral 3’UTRs [31, 78]. The canonical Musashi binding motif, `UAG`, is present at multiple loci within the unstructured repeat of all CHIKV 3’UTR sequences, supporting the biological role of intrinsic lack of structure. Besides protein-aided regulation, RNA interference is a crucial mechanism that controls viral replication in mosquitoes. Host-encoded microRNAs (miRNAs) were shown to interact with the 3’UTRs of Eastern equine encephalitis virus (EEEV) [79] and CHIKV [80, 81]. The miRNA binding site reported by Dubey *et al.* [80] overlaps with the structured Y-shaped element (SL-Y) characterized here. Whether or not the CHIKV genome encodes for endogenous viral miRNAs, as observed in flaviviruses [82], remains to be established.

The structural aspects of alphavirus RNA is understudied when compared to other arboviruses. While repeat motifs were shown to be a crucial architectural pattern in alphavirus UTRs, many studies focused on the role and nature of these sequence repeats, without asking how far RNA structure might be the evolutionary driving force behind UTR evolution. A recent study employing SHAPE structure probing (selective 2’-hydroxyl acylation analyzed by primer extension) experiments could not find evidence for large-scale structural conservation of RNA within alphaviruses [37]. Work in the 5’UTRs of VEEV [83] and SINV [84] could associate RNA structure elements with replication efficiency and antiviral response [85]. We identified structured elements in the 5’UTR of CHIKV, however did not observe strong covariation patterns (data not shown). Likewise, the structured elements SL-a and SL-b in the 3’UTR exhibit only a few covariations. This observed lack of covariation might be due to the high degree of sequence conservation in the 3’UTR. Alternatively, it could also be a consequence of a sampling bias, as many samples in our dataset originate from individual outbreaks and therefore only represent a fraction of the CHIKV population. In contrast to other arboviruses, such as flaviviruses carrying exoribonuclease-resistant RNAs (xrRNAs) [35], the lack of covariations makes it difficult to assert to what extent secondary structure conservation is a driving force of CHIKV UTR evolution. This is underlined by a weak support of structure-associated z-scores for elements SL-a and SL-b (Figure 3 c,d). Contrary, the SL-Y and CSE elements have very stable structures as indicated by strongly negative z-scores. One of the most striking features of the CHIKV 3’UTR is presence of multiple copies of the unstructured UR region separating structured regions. Our findings indicate that the evolutionary forces acting on CHIKV 3’UTR manifest as a process that can rapidly re-arrange structured and unstructured RNA elements, as observed in a recently emerged partial duplication of CHIKV strains isolated from the Caribbean [65]. Consequently, our predictions encourage further experimental validation to assess the structural and functional role of these proposed elements. In particular, it would be highly informative to elucidate whether or not these elements mediate the production of standalone non-coding RNAs. Additionally, experiments could reveal whether these conserved elements act in cis or trans.

To obtain a more comprehensive picture of the 3’UTR phylogeny in the context of diverged CHIKV lineages, we computed a separate phylogeny based exclusively on 3’UTR sequences (Figure S1). Although coding and non-coding parts of the viral genome are subject to different evolutionary pressures, mapping lineage association onto this tree reproduces the topology of the full CDS tree (Figures 1). This is to some extent surprising, as the nature of evolutionary forces acting on both entities is supposed to be different, best illustrated by the requirement to maintain a reading frame. This means that the 3’UTR evolution is not disentangled from CDS evolution in CHIKV, as expected intuitively. On a broader scale, our data provide evidence that CHIKV CDS and 3’UTRs show coherent evolutionary patterns, with leads to the hypothesis that the 3’UTR might represent an evolutionary scaffold that can rapidly replace functional elements, as required for adaptation to environmental change, e.g. by means of recombination [32]. Still, CHIKV 3’UTRs do not exhibit the high degree of plasticity known from other arboviruses with functional 3’UTRs, such as flaviviruses [28, 35].

## Author Contributions

MTW and ABS conceived the study; ABS, RO, RH and MTW analyzed the data; DJ supported the phylogenetic analysis; ILH supported the RNA structure analysis; All authors contributed to writing the paper and approved of the submitted version.

## Acknowledgments

We thank Adiba Hassan, MPH for the valuable discussions during the development of this work. We acknowledge the support of the Department of Bioinformatics and Genomics, The College of Computing and Informatics, and the Graduate School of the University of North Carolina at Charlotte.

© 2019 by the authors. Submitted to *Viruses* for possible open access publication under the terms and conditions of the Creative Commons Attribution (CC BY) license (http://creativecommons.org/licenses/by/4.0/).

