## Supplement for "Updated phylogeny of Chikungunya virus suggests lineage-specific RNA architecture" (from another version of this preprint: version 1, 2019-07-11)

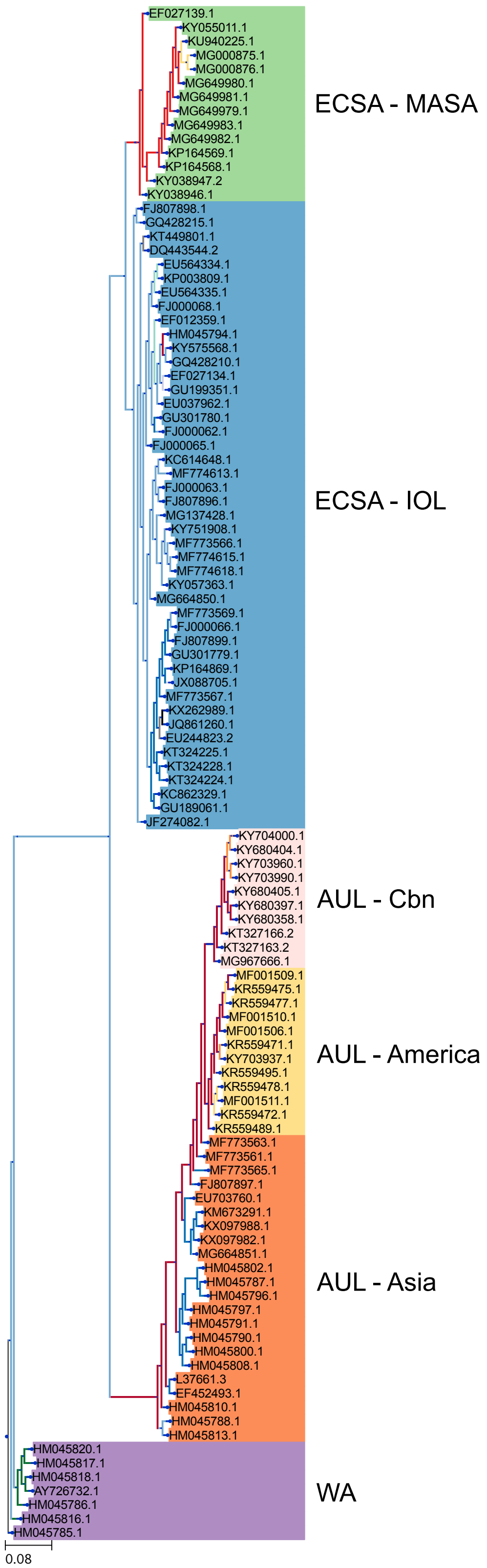


**Fig. 1.** Maximum-Likelihood phylogenetic tree based on a manually curated alignment of 111 complete CHIKV 3'UTRs. Color coding of the tree based on Fig. 1 in main text. Leaves are colored according to lineage, with UTRs harbouring the duplication event in the AUL-American lineage are colored separately.
